# Therapeutic remodeling of the tuberculosis granuloma with 1-methyl-D-tryptophan enhances CD8^+^ T cell-macrophage interactions

**DOI:** 10.1101/2025.10.02.680083

**Authors:** Erin F. McCaffrey, Alea C. Delmastro, Bindu Singh, Annu Devi, Nadia A. Golden, Shabaana A. Khader, Michael Angelo, Deepak Kaushal, Smriti Mehra

## Abstract

Granulomas, the hallmark of tuberculosis (TB) disease, can both restrict *Mycobacterium tuberculosis (Mtb)* dissemination and impede its clearance. Recent studies indicate that indoleamine 2,3-dioxygenase (IDO1), an immunosuppressive metabolic enzyme, limits infiltration of activated T cells and can contribute to TB disease progression. Treatment with 1-methyl-D-tryptophan (D-1MT), a small molecule inhibitor that restores mTOR signaling, has been shown to improve immune responses *Mtb*-infected rhesus macaques. Here, we investigated the impact of D-1MT treatment on TB granuloma architecture using 30-plex high-dimensional issue imaging in rhesus macaques. By spatially mapping 13 distinct cell populations, we found D-1MT treatment corresponded with significantly increased infiltration CD8^+^ T cells into granulomas compared to untreated controls. Notably, these CD8^+^ T cells expressed markers of cell proliferation and cytotoxicity. D-1MT enhanced CD8^+^ T cell infiltration throughout the granuloma, with particularly pronounced effects in the myeloid core, where we observed significantly enhanced spatial interactions between macrophages and CD8^+^, but not CD4^+^ T cells. Our results demonstrate that: (i) effective intra-granulomatous *Mtb* control is associated with the close spatial proximity between CD8^+^ T cells and macrophages, a feature less abundant in uncontrolled pulmonary TB; (ii) IDO1 induction blocks CD8^+^ T cell infiltration and reduces T cell activation and proliferation; and (iii) therapeutic strategies, including D-1MT, that improve intra-granulomatous killing hold strong translational potential.

**Significance statement:** Our understanding of immune mechanisms within the TB granuloma has advanced greatly with the advent of high-resolution single cell multiplexed imaging. Using such imaging, we show that TB granulomas in rhesus macaques, a highly translational model of human TB pathology, are characterized by IDO1-mediated immunoregulation. Early pharmacologic restoration of mTOR signaling via D-1MT treatment can reduce IDO1 enzymatic activity and facilitate the recruitment and function of CD8^+^ T cells within the granulomas. These findings reveal specific mechanisms exploited by *Mtb* to maintain intra-granuloma persistence and underscore immune responses. Future vaccine and therapeutic design should consider these immunoregulatory features to achieve better control of *Mtb* infection.

## Introduction

*Mycobacterium tuberculosis* (*Mtb*) is the leading cause of mortality from infectious disease worldwide, accounting for nearly 1.5 million deaths each year. Relative to other infectious diseases, the reduction in the incidence of tuberculosis (TB) disease over the last 20 years has been unimpressive^1^. This is largely due to the continued lack of a highly efficacious vaccine, lengthy and toxic antimicrobial regimens, and emergence of multidrug resistance. Along these lines, efforts to develop new host-directed therapies for TB will require a deeper understanding of interactions between *Mtb* and the human immune system^2^.

The formation of granulomas is a hallmark of *Mtb* infection. A prototypical granuloma consists of a myeloid-predominant central core region that is highly enriched in infected macrophages and neutrophils and encircled by lymphocytes^3^. These lesions are also characterized by the presence of variable levels of viable *Mtb*, necrosis or cell death, fibrosis, and immune activation^4^. From the perspective of facilitating an effective host response, granulomas play central and seemingly contradictory roles. On one hand, granulomas can sequester *Mtb* and limit dissemination to uninfected tissue sites. On the other hand, upregulated tolerogenic pathways within the myeloid core may limit bacterial clearance, and granuloma-associated inflammation can cause damaging host pathology^5^.

The non-human primate (NHP) model of TB—using experimentally infected rhesus (RM) or cynomolgus (CM) macaques—has been shown to recapitulate the functional diversity of the human TB granuloma^6^. Work in these model systems has demonstrated that granulomas within a single *Mtb-*infected individual can take on a spectrum of fates from complete bacterial clearance to uncontrolled dissemination and inflammation^7–9^. This strongly suggests that local host-bacterial dynamics within an individual granuloma’s microenvironment govern granuloma function, and that granuloma structure and immune cell function are interconnected. These data also suggest that, while some granulomas pose a barrier to controlling *Mtb* infection, others do in fact have the cellular and organizational requirements for bacterial control.

We have previously demonstrated that one of the most highly abundant proteins in both human and NHP TB granulomas is indoleamine 2,3-dioxygenase 1 (IDO1)^5,10^. IDO1 is a metabolic enzyme that catalyzes the conversion of tryptophan to kynurenine^11^. Depletion of tryptophan by IDO1 has been shown to be protective against pathogens, but it also has highly immunosuppressive effects on macrophages and T cells^12,13^. In human TB, IDO1 is expressed by granuloma macrophages alongside other suppressive attributes, such as programmed death ligand 1 (PD-L1), TGFβ, and regulatory T cells (Tregs)^5^. Because we hypothesize IDO1 is suppressing anti-*Mtb* immunity in the granuloma, it is a potential target for host-directed immunotherapy.

One candidate approach for targeting IDO1’s activity is the small molecule 1-methyl-D-tryptophan (D-1MT). D-1MT acts as a mimetic of the amino acid tryptophan, thus reversing the inhibitory impact of tryptophan depletion^14^. Recent work suggests the mechanism of D-1MT is primarily through restoration of mTOR signaling, which is critical for T cell effector functions^14,15^. We have demonstrated that use of this small molecule *in vivo* can reduce IDO1 enzymatic activity in RMs, enhance innate and adaptive immune responses, and reorganize the granuloma to provide greater access to T cells to the granuloma core in RM lungs^10,16^. Given its potential as a host-directed therapy for TB, we sought to more deeply characterize the impact of D-1MT treatment on TB granuloma composition and structure. Here, we used Multiplexed Ion Beam Imaging by Time-of-Flight (MIBI-TOF) to image 30 proteins at subcellular resolution and spatially map 13 cell subsets in TB granulomas from RMs treated with D-1MT or left untreated. Our results first confirm that TB granulomas in both RMs and humans share an immunoregulatory myeloid core characterized by abundant IDO1. Furthermore, we find that D-1MT treatment is associated with a substantial increase in CD8^+^ T cell infiltration and proliferation in the myeloid core. This suggests that inhibition of IDO1 preferentially aids in CD8^+^ T cell immunity in the TB granuloma. Ultimately, this study explores the role of D-1MT as an immunotherapeutic strategy to enhance cellular immunity in granulomas during *Mtb* infection.

## Results

### Application of MIBI-TOF platform to RM tissues and benchmarking with human TB granulomas

We performed an in-depth investigation into the spatial biology of pulmonary TB granulomas in RMs with or without concurrent D-1MT treatment. For these analyses, we curated a cohort of archival formalin-fixed paraffin-embedded (FFPE) lung sections from seven RMs infected with a high dose (∼100-200 CFU) of *Mtb* CDC1551 via the aerosol route that resulted in acute pulmonary TB^10^. Of the seven *Mtb*-infected RMs included in this study, four RMs were untreated (henceforth referred to as controls), and three RMs were treated with D-1MT (45 mg/kg body weight) orally daily, starting one week after *Mtb* infection, as previously reported^10^. All animals were necropsied at weeks 5-8 post-infection, which allows us to characterize early TB granuloma composition and phenotype. In total, our imaging cohort comprised 500 x 500 μm fields-of-view (FOVs), capturing 18 pulmonary granulomas from four control RMs and 15 pulmonary granulomas from three D-1MT-treated RMs (Fig. 1A, Fig. S1).

**Figure 1:**
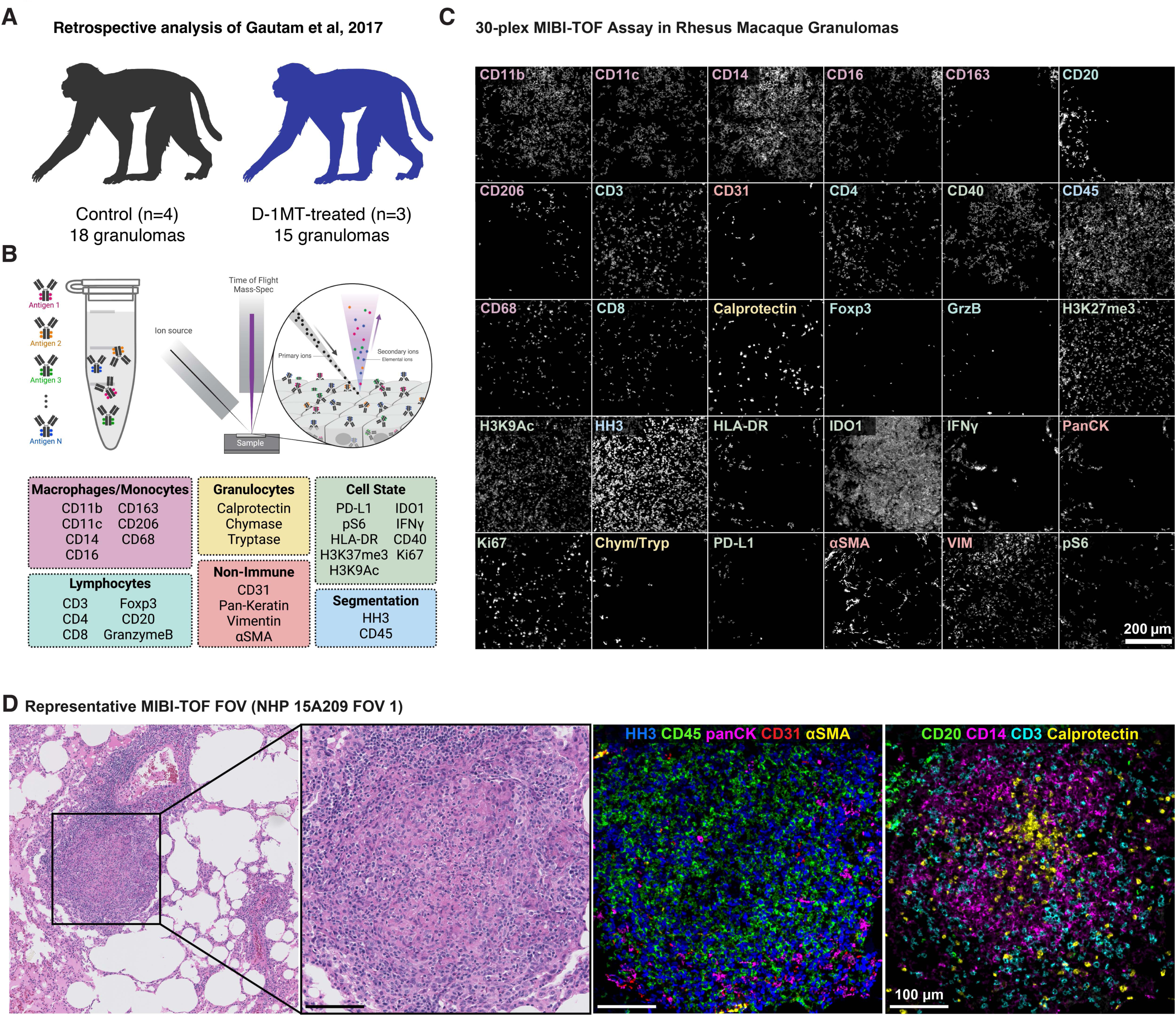
Study design. **(A)** Cohort characteristics, including the number of rhesus macaques (RMs) and the total number of granulomas analyzed by treatment group. Biopsy specimens were obtained from Gautam et. al., 2017. **(B)** Graphical illustration of MIBI-TOF methodology (top) and the list of 31 markers included in the imaging panel (bottom). **(C)** Expression patterns for the 30 marker channels acquired for one representative granulomatous field-of-view (FOV). **(D)** One representative hematoxylin & eosin-stained specimen with inset demonstrating FOV acquired on MIBI-TOF. On the right, two MIBI-TOF overlays, demonstrating major lineage markers: (Third from left) αSMA (yellow), panCK (magenta), CD45 (green), HH3 (blue), and CD31 (red); (Fourth from left) Calprotectin (yellow), CD14 (magenta), CD20 (green), and CD3 (cyan). Scale bars: 100 μm unless otherwise indicated.

All tissues were analyzed with MIBI-TOF, with which we characterized the immunological landscape of TB granulomas from human patients and the CM NHP model of TB^5,17,18^. Prior studies identified the high expression of IDO1 and suggested its immunoregulatory role in anti-*Mtb* T cell responses^9,10,16,19,20^. Therefore, we aimed to perform similar high-dimensional analyses of the composition and phenotype of TB granulomas from D-1MT-treated versus control animals to elucidate the immunoregulatory mechanism of IDO1.

To achieve this, we first generated a 30-plex panel of metal-labeled antibodies cross-reactive with RM epitopes (Fig. 1B-C, Fig. S2). This panel included markers to phenotype major immune and nonimmune cell lineages, including lymphocytes, macrophages, granulocytes, stroma, and epithelium (Fig. 1B-C). Fig. 1D demonstrates the hematoxylin and eosin-stained tissue and two MIBI-TOF overlays of major lineage markers for an example granuloma from one of the D-1MT-treated RMs. We also included markers to evaluate immune regulation (IDO1), cell activation (CD40, HLA-DR, pS6), proliferation (Ki67), cytotoxicity (GranzymeB), Th1 cytokine production (IFNγ), and epigenetic state (H3K37me3, H3K9Ac) (Fig. 1B-C). Representative staining patterns of all the markers included this study are shown in Fig. 1C.

Following low-level processing of the imaging data (see Methods), we performed single cell segmentation with the deep-learning approach, Deepcell^21,22^ (Fig. 2A). Segmented cells were then partitioned into non-immune cells (CD45^−^ panCK^+^/CD31^+^/aSMA^+^) or immune cells (CD45^+^ CD20^+^/CD14^+^/CD3^+^/Calprotectin^+^). Using iterative clustering, non-immune cells were further resolved into fibroblasts, endothelial cells, and epithelial cells, and immune cells were clustered into one of nine subsets (lymphoid: CD4^+^, CD8^+^, regulatory T cells (Tregs), other T cells and B cells; myeloid: macrophages/monocytes, neutrophils and giant cells; and indeterminate: other immune lineage cells) (Fig. 2B). This allowed us to map the single cell composition of each granuloma with our dataset comprising 68,296 cells across the 33 FOVs analyzed (Fig. S3A). A similar study of human TB granulomas previously identified 20 different cell types using 24 different markers^5^. Of these, we find that CD4^+^, CD8^+^ T cells, B cells, neutrophils and giant cells were identified in both human and RM granulomas amongst immune cells while all three stromal cell types (epithelial, endothelial, fibroblasts) were commonly identified (Fig. 2C, Fig. S3B). Unlike the human TB granuloma dataset, we neither performed sub-clustering of macs/monos nor included a marker for γδ T cells, which likely explains the discrepancy between the number of identified subsets. In conclusion, we demonstrate the successful application of the MIBI-TOF platform to RM tissues and bolsters the translational relevance of the NHP TB model for studying human TB disease.

**Figure 2:**
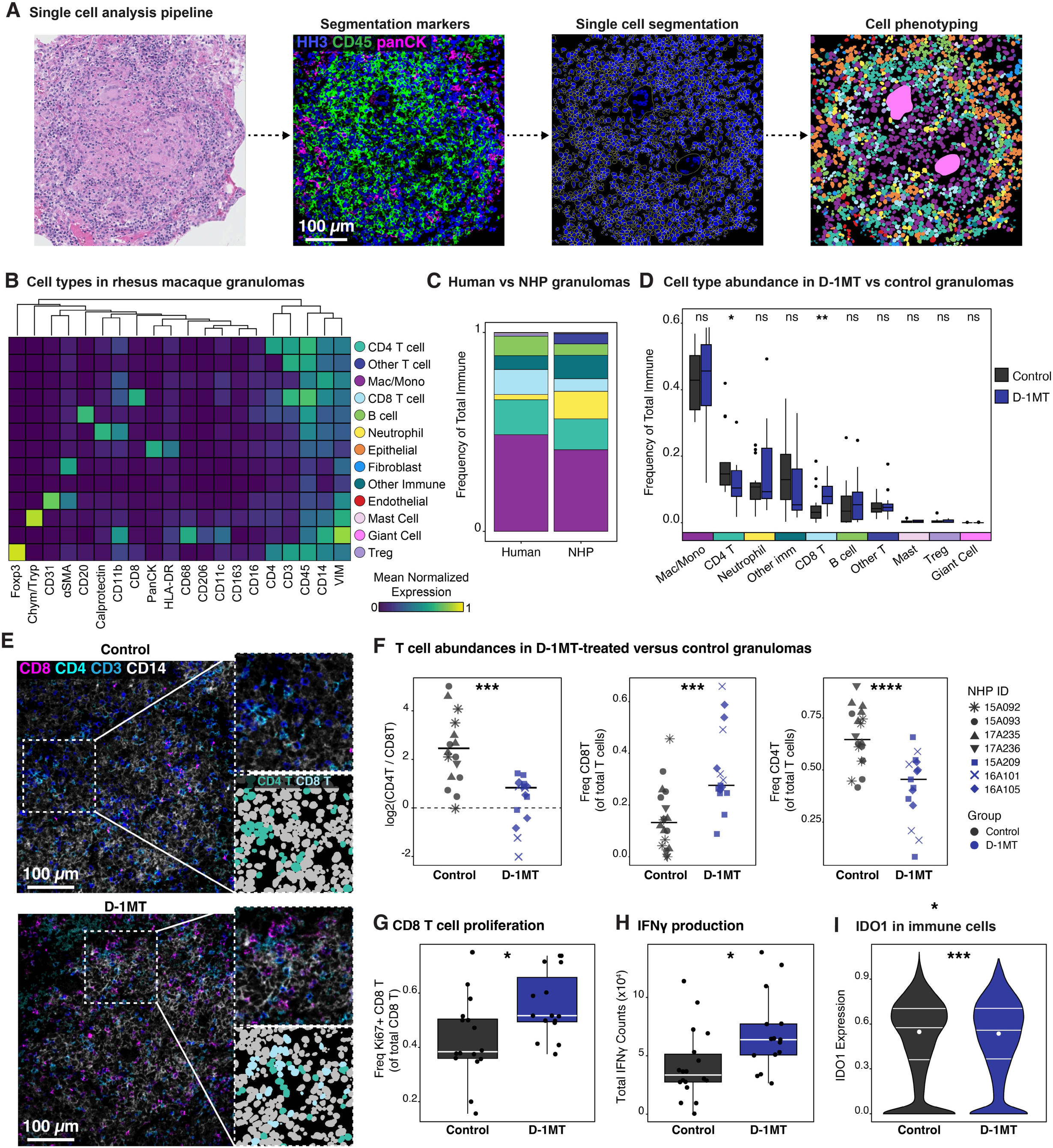
Cellular landscape of rhesus macaque granulomas. **(A)** Graphical overview of the entire analytical pipeline applied to the data set as part of this study, including cell segmentation and cell phenotyping. **(B)** Cell lineage assignments based on mean normalized expression of lineage markers (heatmap columns). Rows and columns are hierarchically clustered (Euclidean distance, average linkage). **(C)** Comparison of immune cell composition of mycobacterium tuberculosis granulomas from humans (from McCaffrey et. al., 2022) and from RMs. **(D)** Frequency distribution comparison of each immune cell subset between granulomas from D-1MT-treated RMs and control RMs. **(E)** MIBI-TOF expression overlays (top – control RMs and bottom – D-1MT-treatment RMs) demonstrating T cell subsets: CD8 (magenta), CD4 (cyan), CD3 (blue), and CD14 (white). Smaller insets emphasize expression patterns (top) and illustrate the presence of CD4+ T cells and CD8+ T cells in the cellular phenotype map (CPM): CD4+ T cells in teal and CD8+ T cells in light blue. **(F)** Comparison of the CD8+ T (center) and CD4+ T (right) cell frequencies of total T cells between control and D-1MT-treated RMs, on a per-granulomatous FOV basis. (Left) Comparison of the log2 ratio of CD4+ T and CD8+ T frequency between control and D-1MT-treated RMs. **(G)** Frequency of Ki67+ CD8+ T cells among control and D1-MT-treated RMs. **(H)** Total IFN*γ* counts within granulomatous FOVs, compared between control and D-1MT-treated granulomatous lesions. **(I)** Violin plots denoting IDO1 expression among all immune cells, stratified by treatment group. Scale bars: 100 μm. All p-values were calculated with a Wilcoxon Rank Sum test (*p < 0.05, **p < 0.01, ***p < 0.001, ****p < 0.0001).

### Comparison of cellular composition and state in D-1MT-treated versus control TB granulomas

We next sought to compare the cellular composition of TB granulomas from D-1MT treated versus control RMs by analyzing both the count and frequency of all cell subsets in each group (Fig. 2D, Fig. S3C). Of all the subsets identified, we found that only CD8^+^ T cells and CD4^+^ T cells varied between groups with a significant increase CD8^+^ T cell frequency (of total immune cells, p = 0.0036) and count (p = 0.0045) in the D-1MT-treated animals (Fig. 2D, Fig. S4A-B). Taking a closer look at the lymphocyte compartment of the granulomas, we found that this increase in CD8^+^ T cell frequency (of total T cells, p = 0.00063) coincided with a decrease the proportion of CD4^+^ T cells (p = 9.2e-5), leading to granulomas that were more CD8^+^ T cell-skewed (Fig. 2E-F). In our prior study of these animals, we previously observed increases in both CD4^+^ and CD8^+^ T cells in the lung and bronchoalveolar lavage (BAL) fluid of D-1MT-treated RMs, yet, in the granuloma, it was only the CD8^+^ T cells that were elevated in the D-1MT-treated RMs. This is notable deviation from the CD4^+^ T cell dominated response typically associated with anti-*Mtb* immunity in the granuloma.

We next asked whether this shift in the T cell composition of granulomas from D-1MT-treated RMs corresponded with changes in cell state or function. We found that CD8^+^ T cells, but not CD4^+^ T cells, displayed elevated levels of Ki67, a marker of cellular proliferation (p = 0.02, Fig. 2G, Fig. S4C), suggesting that increased CD8^+^ T cell abundance in the setting of D-1MT results from T cell expansion in the granuloma itself as opposed to increased recruitment. We found no difference in the proportion of CD8^+^ T cells that express GranzymeB between D-1MT-treated and control RMs (Fig. S4D), indicating no difference in the cytotoxic capacity of CD8^+^ T cells with treatment. However, we did find increased quantities of IFNγ in granulomas from D-1MT-treated RMs relative to those from control RMs (p = 0.02, Fig. 2H). Lastly, while a reduction in IDO1 enzymatic activity was reported by us in these animals (as indicated by the tryptophan: kynurenine ratio; data not shown), we found only modest, yet significant, reduction in the expression of IDO1 across immune cells in bulk (p = 0.00098, Fig. 2I) but not in macs/monos alone (Fig. S4E). We found no significant difference in the presence of any other functional marker (CD40, HLA-DR, CD40, pS6, H3K9Ac, and H3K27me3) at either the single-cell or image level (data not shown). Overall, D-1MT treatment is associated with an expansion of CD8^+^ T cells in the granuloma and a switch to a more Th1-like cytokine environment.

### D-1MT treatment restructures the spatial organization of TB granulomas

T cell–macrophage interactions are crucial for mediating host immunity and improving mycobacterial killing. Canonically, the myeloid core is depleted of effector CD4^+^ and CD8^+^ T cells and instead is preferentially infiltrated by proliferating Tregs. In previous work using conventional fluorescence microscopy, we previously observed that granulomas from D-1MT-treated animals exhibited increased T cell infiltration^23^. Based on the observed elevation of CD8^+^ T cell frequency and proliferation, we hypothesized that D-1MT treatment—by restoring mTOR signaling and inhibiting IDO1—could promote CD8^+^ T cells infiltration into the myeloid core.

To test this, we automatically defined and masked the myeloid core region of each granuloma (Fig. 3A). Cells within these regions were assigned as ‘myeloid core-infiltrating,’ while those outside the mask were assigned to the lymphocytic cuff (Lcuff)/periphery (Fig. S5A). We found that CD8^+^ T cells were significantly enriched in the myeloid core of granulomas from D1MT-treated RMs relative to control RMs (p = 0.0045, Fig 3B). The CD8^+^ T cell frequencies were also significantly higher when measured in the Lcuff/peripheral region of granulomas, but to a lesser extent than in the myeloid core (p = 0.027, Fig 3B).

**Figure 3:**
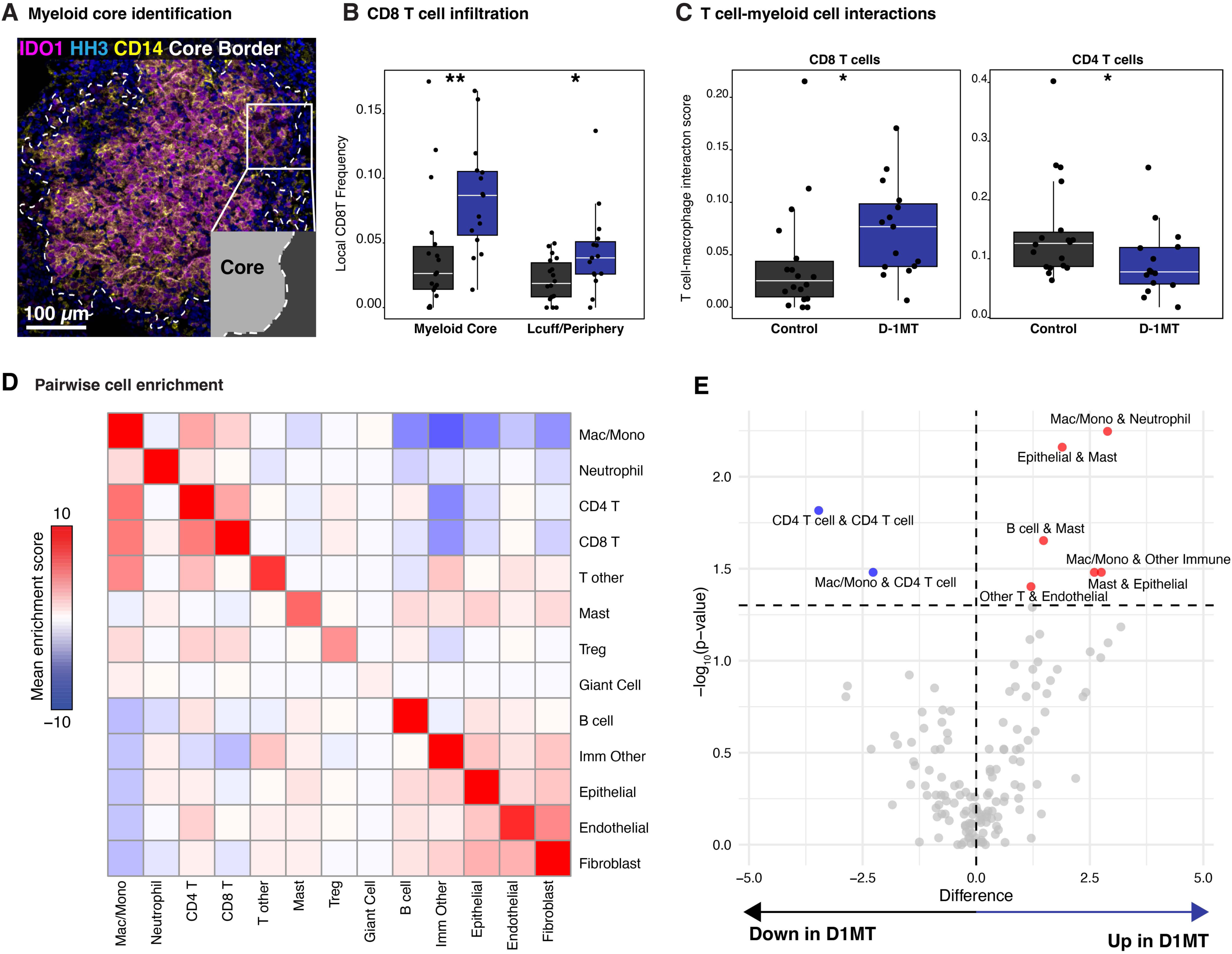
CD8 T cell-macrophage/monocyte interactions are more prevalent in D-1MT-treated rhesus macaque granulomas. **(A)** Representative granuloma with myeloid core border annotated (white, IDO1= magenta, CD14 = yellow, HH3 = blue) with zoomed inset annotated the core (light grey) and periphery (dark grey). **(B)** Quantification of the local frequency of CD8+ T cells of total cells in the myeloid core or lymphocytic cuff (Lcuff)/periphery. **(C)** CD4+ T cell-myeloid cell (right) and CD8+ T cell-myeloid cell (left) mixing scores in control versus D-1MT-treated granulomas. **(D)** Pairwise cell subtype enrichment (or depletion) in all granulomas studied, and **(E)** the effect of D-1MT treatment, shown as a volcano plot. All p-values were calculated with a Wilcoxon Rank Sum test (*p < 0.05, **p < 0.01).

We next assessed T cell-macrophage spatial associations to determine if D-1MT treatment increased cellular interactions between lymphocytes and myeloid cells. For quantification, we employed a mixing score that quantified the number of CD4^+^ or CD8^+^ T cells interactions with macrophages/monocyte relative to the number of homotypic macrophage/monocyte interaction^18,24^. This analysis revealed that the interactions between CD8^+^ T cells and monocyte-lineage cells were indeed significantly increased in D1MT-treated RMs relative to control RMs (p = 0.011, Fig 3C). Interestingly, granulomas from D-1MT-treated RMs had lower interaction scores between CD4+ T cells and macrophages/monocytes compared to those from control RMs (p = 0.018, Fig 3C), suggesting that increased CD8^+^ T cell interactions may come at the expense of macrophage/monocyte-interactions with CD4^+^ T cells.

Given the increased infiltration of the granuloma myeloid core by CD8^+^ T cells after D-1MT treatment, we next conducted pairwise cell enrichment to better understand the spatial patterning of all cell subsets^5,24^. Consistent with the compartmentalization of macrophages in the myeloid core, we found that macrophages/monocytes were spatially enriched with themselves (Fig 3D). Furthermore, T cells (CD4^+^ T cells, CD8^+^ T cells, and other T cells) displayed the strongest spatial enrichment with other T cells, consistent with their predominance in the lymphocytic cuff. When comparing granulomas from D-1MT-treated RMs with those from control RMs, we observed spatial enrichment of macrophages/monocytes with neutrophils and other immune cells as well as mast cells with B cells and epithelial cells in granulomas from D-1MT-treated animals (Fig. 3E, Fig. S5B). Conversely, spatial enrichment of CD4^+^ T cells with other CD4^+^ T cells and with macrophages/monocytes were significantly reduced with D-1MT treatment (Fig. 3E). Taken together, D-1MT treatment is associated with spatial remodeling of the granuloma that favors increased interactions between CD8^+^ T cells and macrophages.

### Validation of increased CD8^+^ T cell infiltration into the granulomas of D-1MT-treated animals by confocal microscopy

To independently validate our MIBI-TOF findings, we stained FFPE sections from the same granulomas analyzed via MIBI-TOF with antibodies specific for CD8 and Granzyme B. We then imaged those samples with confocal microscopy (Fig. 4A-B). Quantification of CD8^+^ T cell density across multiple sections revealed a significant increase in CD8^+^ T cells per unit area within granulomas from D-1MT-treated RMs (mean = 8.92%) compared to untreated controls (mean = 1.34 %) RMs (p=0.0006, Fig. 4C). Similarly, we observed that levels of Granzyme B expression (both on CD8^+^ T cells and on total CD3^+^ T cells) was comparable in treated versus untreated animals (Fig 4C). These results independently validate our high-dimensional imaging findings and support the conclusion that D-1MT treatment enhances CD8^+^ T cell infiltration into TB granulomas.

**Figure 4:**
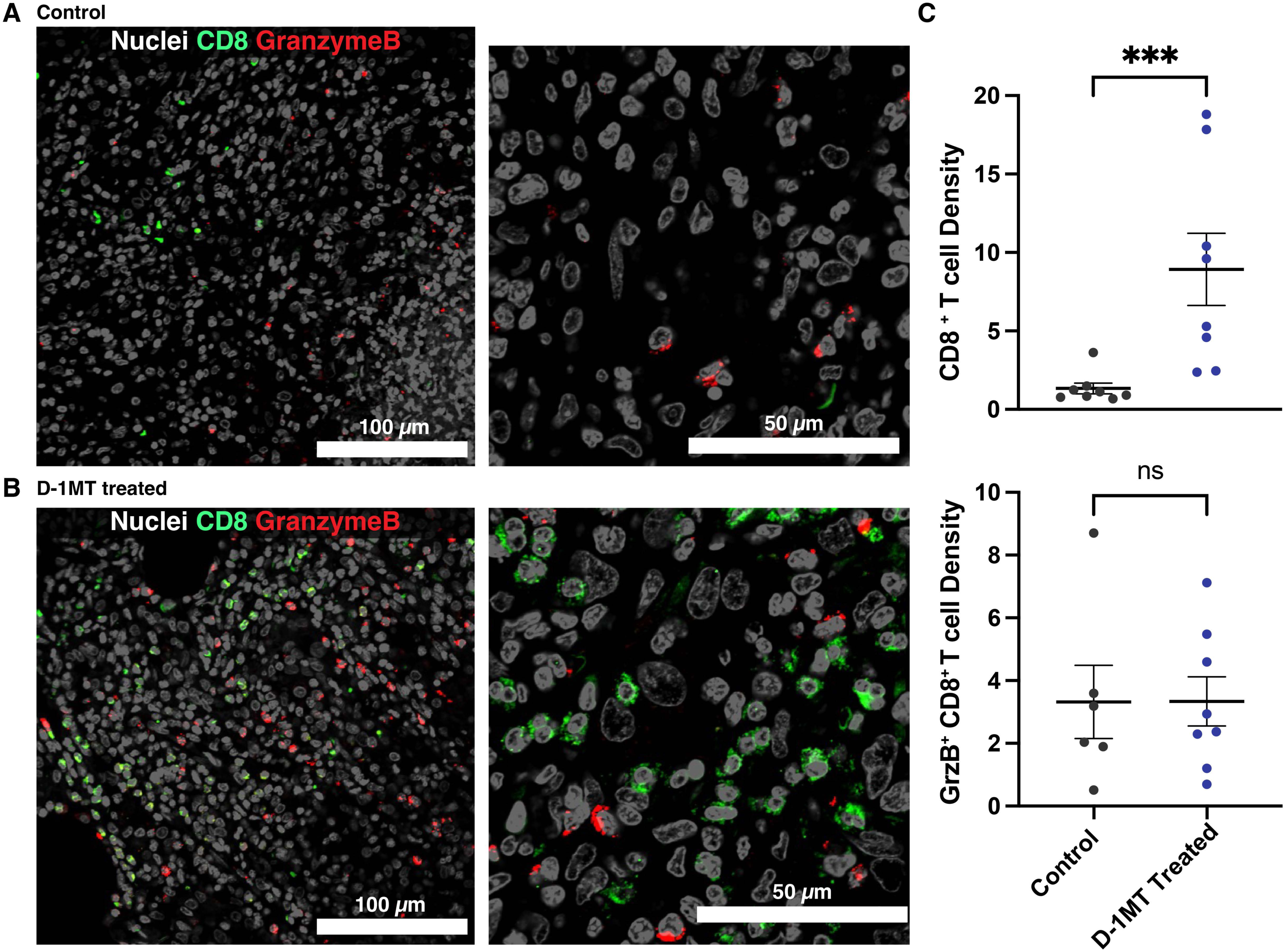
Confocal microscopy validates increased CD8+ T cell infiltration into the granulomas of D-1MT-treated rhesus macaques. CD8 (green), GranzymeB (red), and nuclear staining (white) in *Mtb*-infected **(A)** untreated RM granulomas (20x left; 63x right) and **(B)** D1MT-treated RM granulomas (20x left; 63x right). **(C)** Frequency of CD8+ T cells (of all cells), stratified by granulomatous lesions from control RMs and D-1MT-treated RMs. **(D)** Percent of GranzymeB (GrzB) positivity among CD8+ T cells, stratified by granulomatous lesions from control RMs and D-1MT-treated RMs. All p-values were calculated with a Student’s t-test (two tailed) (***p < 0.001).

## Discussion

Granulomas are critical for the immune control of *Mtb* infection; however, not all granulomas effectively control TB disease. TB granulomas display heterogeneity in their primary function – *Mtb* killing capacity – as well as in cellular immune responses, during both the latent control of *Mtb* infection (LTBI) and progression to active TB disease^7,25,26^. The basis of this functional heterogeneity is unclear. RMs are a robust, pre-clinical model for studying TB and can recapitulate features of both active TB and LTBI. Importantly, RM granulomas mirror the morphology and physiology observed in human disease^27^. Immune correlates of TB disease in the lungs of RMs overlap significantly with those observed in the blood of human TB progressors^28^. The accumulation of myeloid cells (e.g., plasmacytoid dendritic cells, pDCs, that express type I interferon (IFN); inflammatory, IDO1^+^ interstitial macrophages, IMs; and myeloid derived suppressor cells, MDSCs) is a dominant feature of granulomas during TB disease in both humans and RMs^16,29–31^. A protective role for lymphoid features, including increased inducible bronchus-associated lymphoid tissue and cytolytic effector-expressing NK cells, has also been defined within granulomas during *Mtb* control^29,32,33^. The role of type I IFN instead is controversial, with some studies indicating a pathological effect and others suggesting a protective role^34–36^. Type I IFN signaling, which is orchestrated by pDCs and IDO1 signaling on IMs, likely shapes the efficacy of individual granulomas in controlling *Mtb*. Collectively, there are myriad cellular players and pathways influencing granuloma function during TB.

This study utilized MIBI-TOF on preserved lung granuloma sections from experimentally *Mtb*-infected RMs to better understand granuloma structure and function, including the role of IDO1 in suppressing anti-TB responses. We have recently used this innovative technique to spatially profile both human and CM TB granulomas at a single cell resolution^5,18^. Granulomas from human TB patients were characterized by an intense immunosuppressive phenotype with high IDO1 and PD-L1 expresssion^5,18^. In the current study, we found that most immune and stromal cell populations present in human TB granulomas are also detected in RM TB granulomas. We also determined that immune cells make up at least 75% of all the cells in the RM granulomas. Approximately half of all the immune cells detected in RM TB granulomas are macrophages or monocytes. Besides neutrophils, which make up about 10% of immune cells, other myeloid cell populations, such as mast cells and giant cells, were detected at much lower frequencies. Amongst lymphocytes, CD4^+^ T cells were the most frequent followed by B cells, CD8^+^ T cells, other (likely γδ T cells) T cells, and Tregs, in order of decreasing frequency.

The finding that CD8^+^ T cell frequency is significantly increased in granulomas, specifically in the myeloid core, of D-1MT-treated macaques is of interest. We have earlier shown that D-1MT treatment results in the remodeling of the TB granuloma with greater T cell access to the granuloma core, but their phenotyping as mostly CD8^+^ T cells informs us about their potential role in limiting *Mtb* intra-granulomatously. The role of CD8^+^ T cells in controlling *Mtb* infection is not fully understood. Thus, while CD8^+^ T cells were earlier thought to be unimportant in the immune control of *Mtb*, their role is now being increasingly recognized as important yet complex^37^. Mice lacking the ability to generate functional CD8^+^ T cells or those where these cells were depleted have an impaired ability to control *Mtb*^38–40^. CD8^+^ T cells recruited in response to *Mtb* infection are antigen-specific, exhibit memory phenotype, and impart protection independent from CD4^+^ T cells during vaccination^41–44^. A role for CD8^+^ T cells in anti-TB immunity in macaques is known^45^. According to our results, activated CD8^+^ T cells gain access to the granuloma core in the absence of IDO1 activity and restoration of mTOR signaling. These CD8^+^ T cells are highly proliferative (Ki67^+^), associate with a Th1 cytokine shift (increased IFNγ and decreased IDO1), express cytolytic effectors (Granzyme B). Thus, inhibition of IDO1 via D-1MT remodels the TB granuloma, providing activated, proliferative, cytokine producing CD8^+^ T cells access to the lesion core in the vicinity of infected myeloid cells. These results are supported by the evidence that overexpression of IDO1 attenuates the generation of central memory and effector CD8^+^ T cells, while suppressing IDO1 activity promotes their generation^46^, accompanied by defects in production of granule cytotoxic proteins, perforin, and Granzyme A/B^47^. Furthermore, CD8^+^, and not CD4^+^, T cells dominate the immunosuppressed milieu in response to mycobacterial infection^48^. We hypothesize that the greater interaction between these CD8^+^ T cells and infected myeloid cells leads to enhanced *Mtb* killing, resulting in better control of infection. A strong rationale therefore exists to understand the role, mechanistically, of CD8^+^ T cells in the killing of *Mtb* intra-granulomatously.

## Methods

### Nonhuman primate samples, antibody preparation and tissue staining for MIBI-TOF

Rhesus macaques (RMs) were infected and treated as described by Gautam et al^10^. 14 formalin-fixed paraffin-embedded (FFPE) blocks of pulmonary tissues from four control, *Mtb* CDC1551-infected and three *Mtb* CDC1551-infected/D-1MT treated RMs were used for MIBI-TOF imaging experiments. In total, 18 granulomas from controls and 15 from D-1MT-treated RMs were analyzed (Fig. 1A). 5 μm serial sections of each specimen were stained with hematoxylin and eosin and two-six 500 μm^2^ FOVs were selected from each block for imaging via MIBI-TOF. To accurately capture granuloma microanatomy in these regions, we prioritized imaging FOVs with smaller, non-necrotizing cellular granulomas. One RM lymph node and one RM spleen specimen were included as technical controls. Antibodies were conjugated to isotopic metal reporters, diluted, stored, reconstituted and used as described previously^5^. Information on the antibodies and metal reporters used for NHP experiments and staining concentrations is in Table S1.

### MIBI-TOF Imaging

Imaging was performed using a MIBI-TOF instrument with a Hyperion ion source as previously described^17^. All samples were stained with a 30-plex antibody panel (Table S1, Fig. 1B-C) and imaged with the MIBI-TOF platform. Multiplexed image sets were extracted, slide background-subtracted, denoised, and aggregate filtered using a custom low-level processing pipeline described by us previously^49^. In addition to these processing steps, image compensation was performed to account for signal spillover due to adducts and oxides for the following interferences: Collagen-1 to IDO1 and Lag3, H3K9Ac to panCK and MPO, Chym/Tryp to MPO, Ki67 to CD209, CD20 to CD16, CD16 to IFNγ, CD11c to IDO1, and HLA-DR-DQ-DP to CD11b.

### Cell Segmentation and Phenotyping

Nuclear segmentation was performed using an adapted version of Deepcell, a convolutional neural network that can be trained to predict single-cell segmentation across a range of biological platforms^21,22^. This algorithm was trained on 2600 images of cells of diverse shapes and morphologies from nine different tissue types. Rather than predicting the nucleus and performing a radial expansion, this algorithm directly predicts the shape of the entire cell. The updated algorithm was used to generate segmentation predictions for each cell in the image with HH3 and CD45/pan-Keratin as nuclear and cell membrane channels, respectively, as input. Multinucleated giant cells were manually segmented in ImageJ and merged with the Deepcell-generated segmentation masks. Single cell data was extracted and normalized as described above. Single cell data was extracted for all cell objects and area-normalized. Cells with a sum of less than 0.1 area-normalized counts across all lineage channels were excluded from analysis. Single cell data was linearly scaled with a scaling factor of 100 and asinh-transformed with a co-factor of 5. All mass channels were scaled to 99.9^th^ percentile. To assign each cell to a lineage, the FlowSOM clustering algorithm was used with the Bioconductor “FlowSOM” package in R. FlowSOM clustering was applied to assign each segmented cell to one of thirteen phenotypes (Fig. 1B), and the proportion of cell types was quantified per granuloma FOV. Marker thresholds for each channel were automatically assigned using the MetaCyto silhouette scanning approach^50^.

### Spatial Statistics

To compare spatial features between groups, the IDO1 channel was used to produce a mask of the myeloid core as its expression appeared highly compartmentalized to regions expressing CD11c, CD11b, and CD14. The mask was produced by first capping (cap = 10 counts), Gaussian-blurring (sigma = 5), and binarizing the IDO1 channel. Next, close objects in the mask were connected using Matlab’s ‘imclose’ function and objects with a size less than 10000 pixels were filtered out of the mask. The mask was further smoothed by filling in holes with the ‘imfill’ function, dilating the mask, and applying active contouring. Anything within the mask boundary was considered part of the myeloid core, while anything outside the mask was annotated as part of the ‘periphery.’ The local proportion of CD8^+^ T cells was quantified in both zones relative to the total number of cells in each zone. To further assess the interaction between lymphocytes and myeloid cells a mixing-score was calculated to quantify the degree of interaction between CD4^+^ or CD8^+^ T cells with macrophages and monocytes. The score was adapted from a tumor-immune mixing score presented by Keren et. al. 2018 and was calculated as: (total # CD4^+^ or CD8^+^ T cell-myelomonocytic interactions / total # myelomonocytic-myelomonocytic interactions), where an interaction was defined as two cells with < 10 μm centroid-centroid distance^24^. The higher the score, the higher degree of T cell-myelomonocytic cell interactions. To characterize cell-cell interactions, pairwise enrichment analysis was applied (adapted from Keren et. al. 2018). For each cell, the physical distance to all other cells in the FOV was calculated and stored as a distance matrix. For each cell type A and B, interactions within 100 pixels (∼50μm) were counted as close. Bootstrapping was applied to evaluate whether the number of close interactions is significant compared to interactions when the location of cell type B was randomized. This process was repeated 1000 times to generate a null distribution, and a z-score was calculated to assess the deviation of the actual number from the null distribution. The z-scores were averaged across all FOVs to generate a heatmap of the pairwise enrichment, with the rows hierarchically clustered. The z-scores per FOV for each cell-cell interactions were sorted into control and D-1MT groups, with which Wilcox hypothesis testing was performed, and the log2 fold change was calculated.

### Software

Image processing was conducted with Matlab 2016a and Matlab 2019b. Statistical analysis was conducted in Matlab 2016a, Matlab 2019b, and R version 3.6.2. Data visualization and plots were generated in R. Representative images were processed in Adobe Photoshop and figures were prepared in Adobe Illustrator. Schematic visualizations were produced with Biorender.

### Confocal Microscopy

To validate the findings of lung granuloma MIBI-TOF imaging, multilabel immunohistochemistry was performed on *Mtb*-infected RM lungs with active TB with and without D-1MT treatment. The lung sections were stained for anti-CD8 antibody (Polyclonal, Cat no: HPA037756, Sigma-Aldrich) and anti-Granzyme-B antibody (Clone-GrB-7, Cat no: M7235, Dako). DAPI was used for nuclear staining. Images were captured using Zeiss LSM-800 confocal microscope at 20X and 63X magnification. For quantification, the slides were scanned on Zeiss Axio Scan Z1, and CD8^+^ T cells as well as Granzyme B expression in the granuloma regions of the lung were quantified using HALO software (Indica Labs).’

## Supporting information

All supplemental tables

## Data availability

The MIBI-TOF dataset analyzed here is available through Mendeley’s data repository at DOI: 10.17632/x5sf8gpr67.1.

## Code availability

All original code utilized in this study can be access at https://github.com/angelolab/publications/tree/main/2025-McCaffrey-Delmastro_etal_D1MT.

## Author Contributions

E. F. M., A. C. D., S.M, B. S., A. D. and N. A. G., conducted research; E. F. M, A. C. D., B. S., M. A. and S. M., analyzed data; S. M., M. A., and D. K. provided funding for research; S. M., E. F. M., A. C. D., and W.R.J wrote the paper; D. K, M. A., and S. A. K. helped with the interpretation of the data and edited subsequent drafts.

## Acknowledgments

This research was supported by NIH grants AI134245, AI181701 and AI128130 to SM, AI111914, AI134240, AI138587 and AI184581 to DK and by institutional grants OD010442 and OD028732. E.F.M. was supported by the National Science Foundation (graduate research fellowship grant 2017242837) and training grant 5T32AI007290. M.A. was supported by the National Institutes of Health (grants CA20997105, OD01982205, CA24063801A1, AG06827902, CA24663303, CA22952904, CA22430901, AG05791504 and AG05628705), the Department of Defense (contracts W81XWH2110143), the Welcome Trust and other funding from the Bill and Melinda Gates Foundation, Cancer Research Institute, the Parker Center for Cancer Immunotherapy and the Breast Cancer Research Foundation. E.F.M. is supported by the Division of Intramural Research, NIAID/NIH. A.D., B.S., D.K., and S.M. are also supported by Tuberculosis Research Advancement Center (AI168439); A.D., S.M., and D.K. are also supported by the Texas Developmental Center for AIDS Research (AI161943). A.D. and B.S. are also supported by a Texas Biomedical Research Institute Forum Grant. SAK is Bernard and Betty Roizman Professor in the Department of Microbiology, University of Chicago College of Medicine. This research was supported in part by the Intramural Research Program of the National Institutes of Health (NIH). The contributions of the NIH author were made as part of their official duties as NIH federal employees, are in compliance with agency policy requirements, and are considered Works of the United States Government. However, the findings and conclusions presented in this paper are those of the author and do not necessarily reflect the views of the NIH or the U.S. Department of Health and Human Services.

---

**Figure S1:**
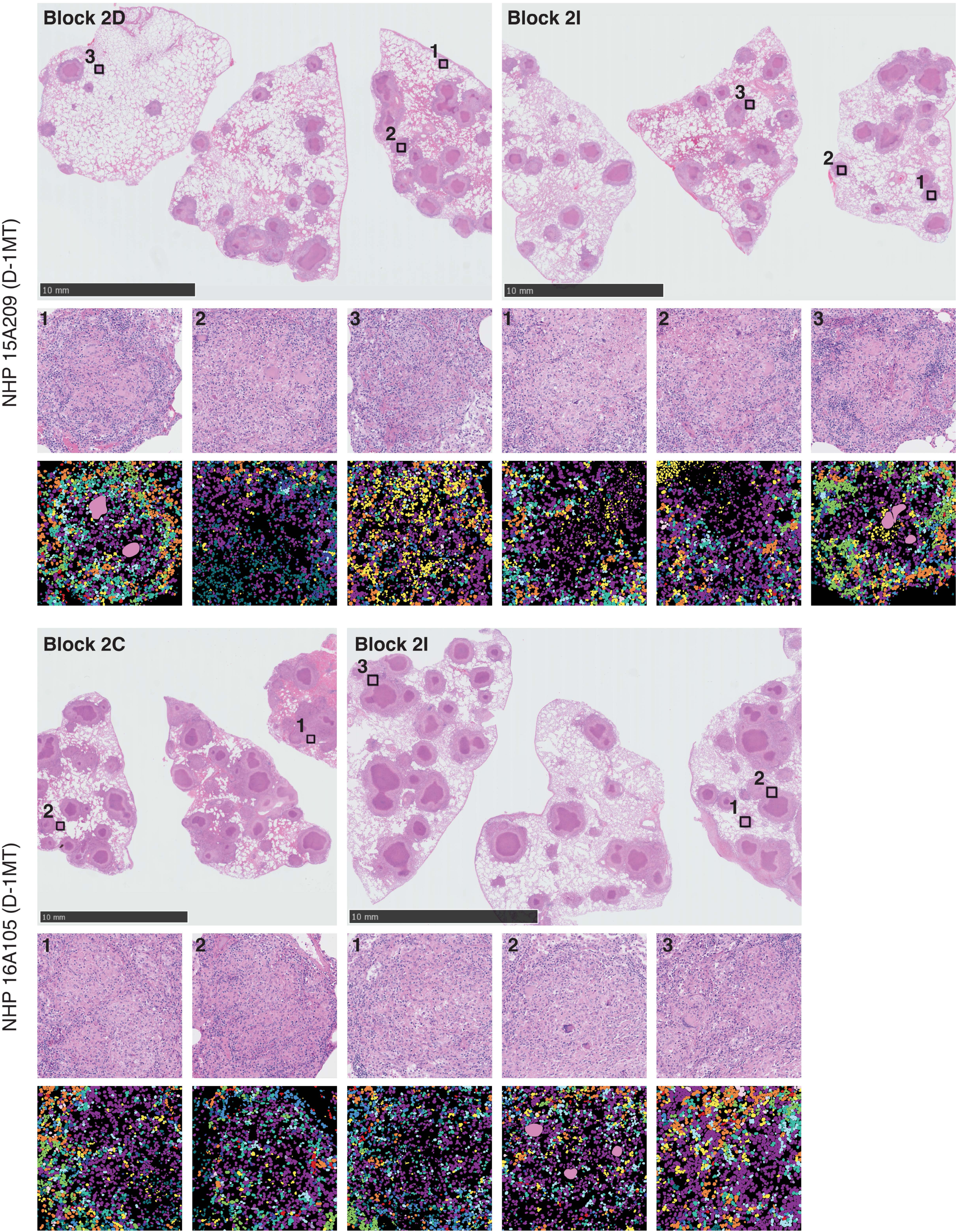
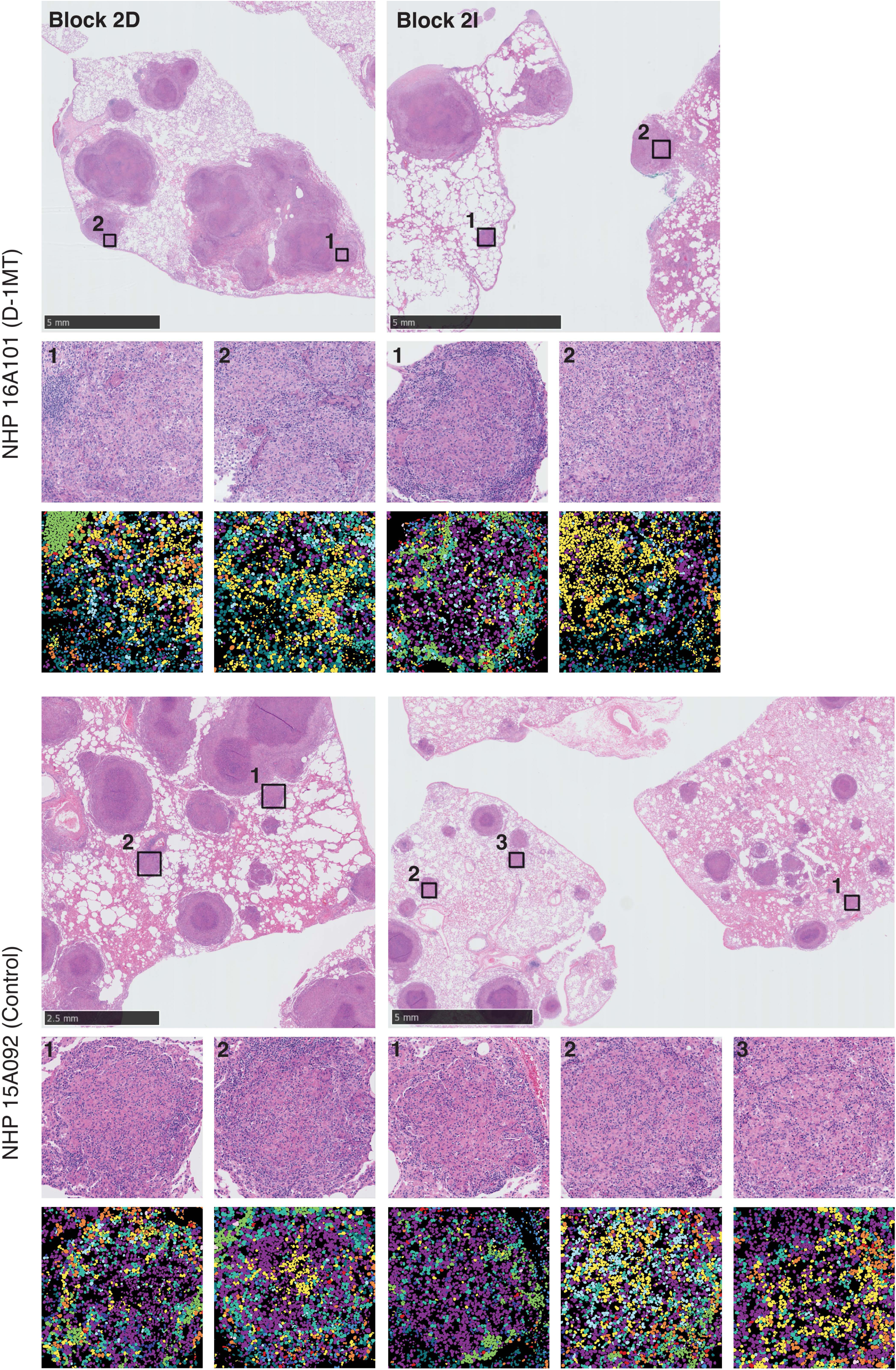
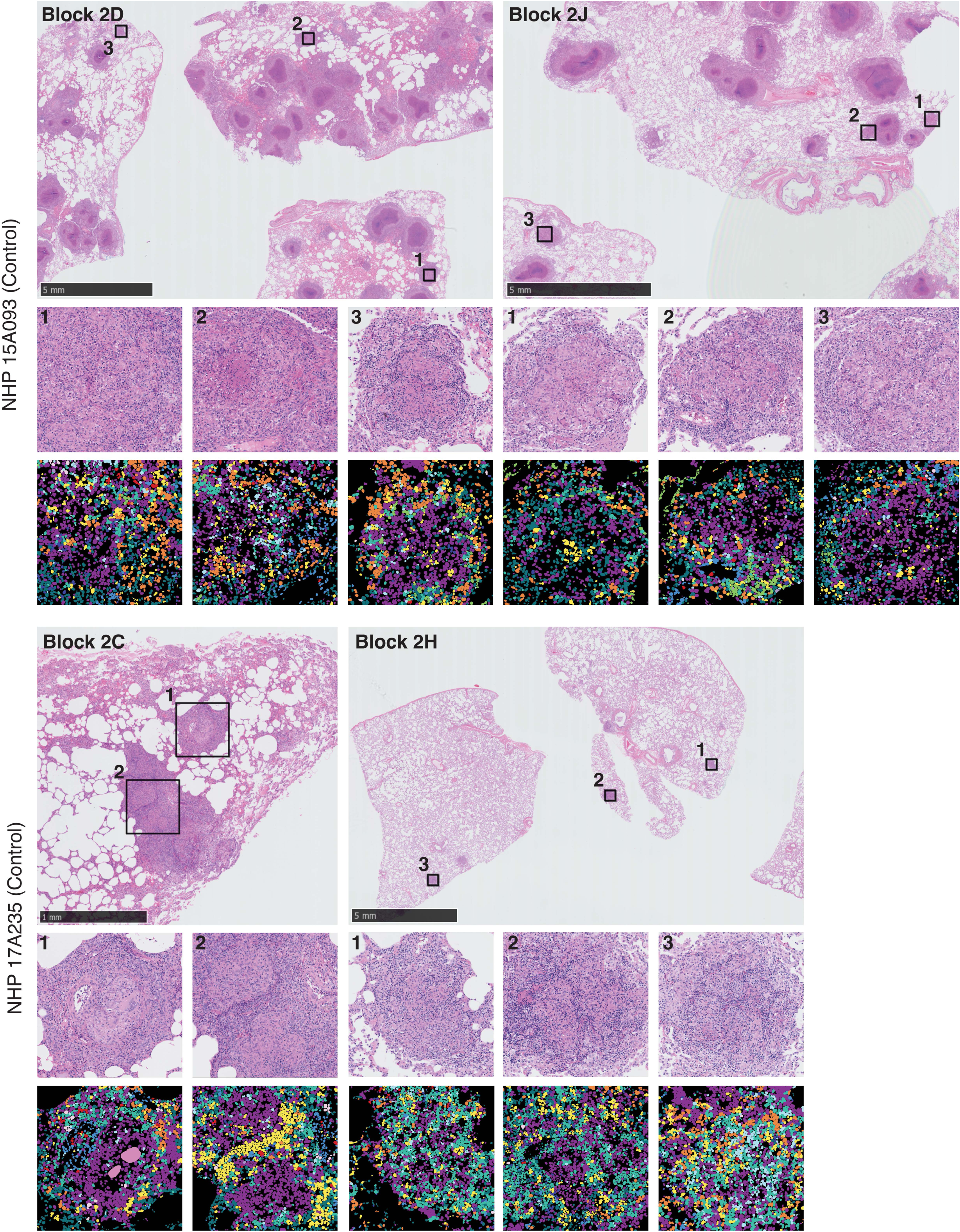
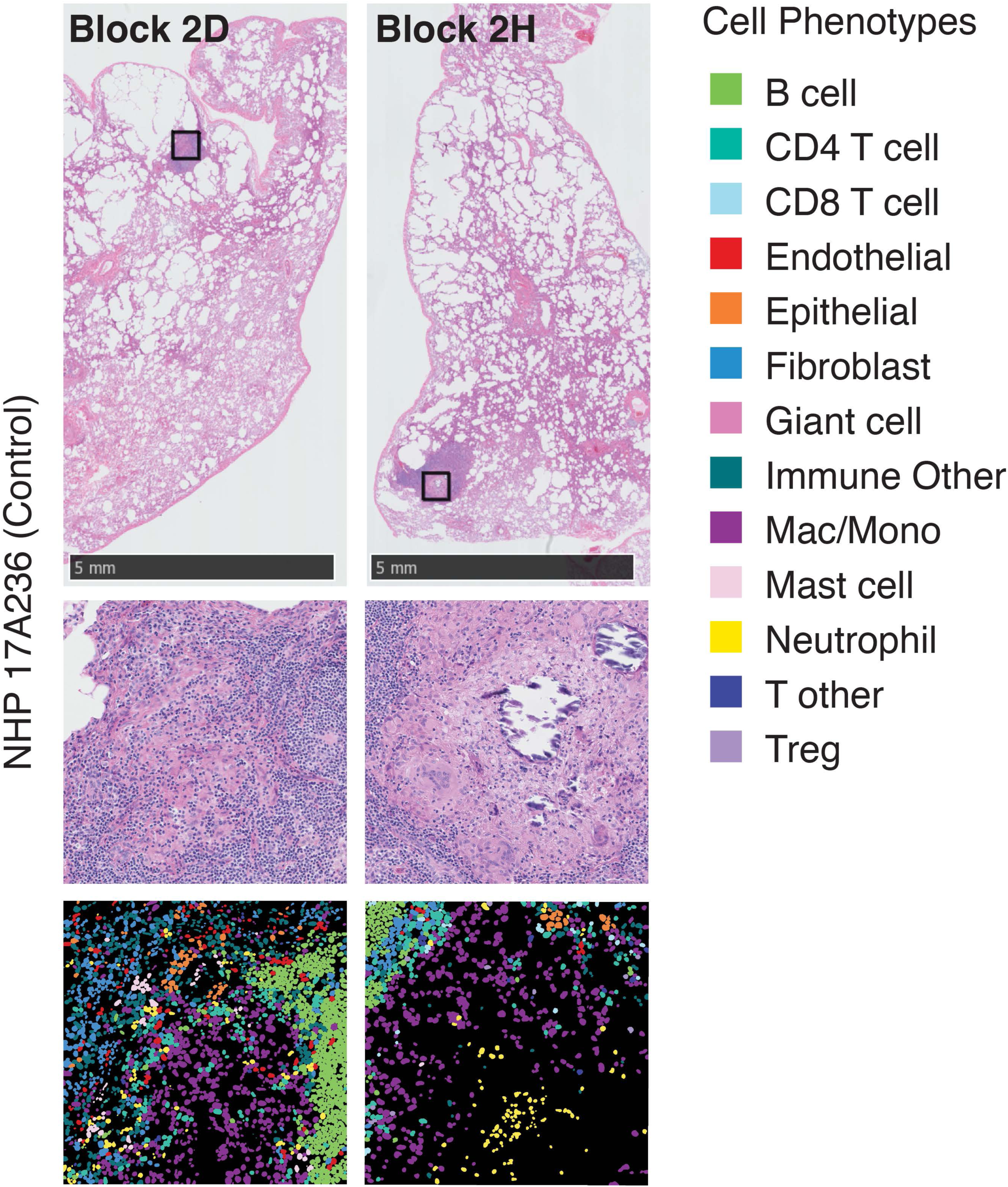
FOV selection and acquisition for each FFPE specimen by rhesus macaque. For each non-human primate (NHP), the first row demonstrates the hematoxylin and eosin-stain of the FFPE specimen, which the locations of each FOV. The second row includes the insets for each 500 x 500 μm^2^ FOV acquired. The third row has the cell phenotype maps (CPMs) for each FOV acquired with MIBI-TOF.

**Figure S2:**
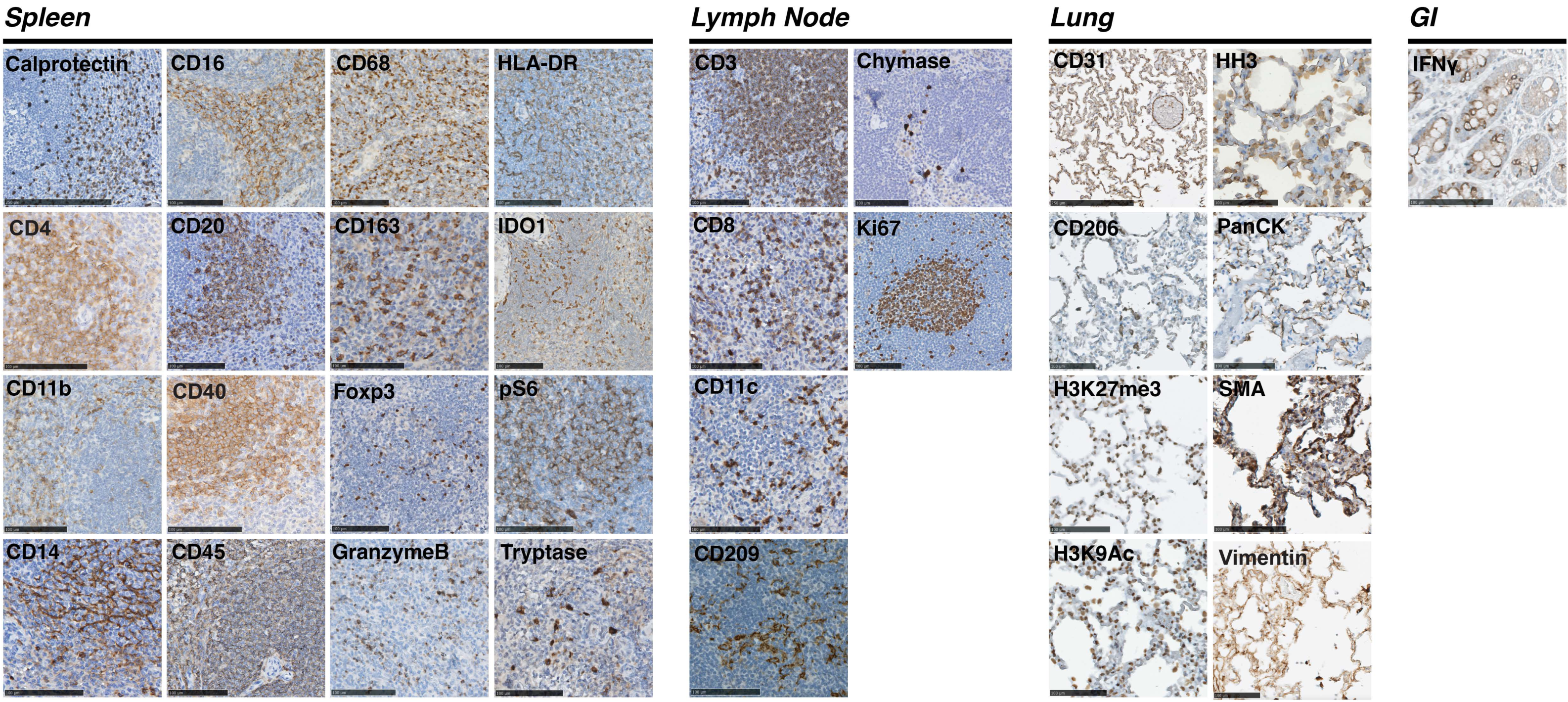
Validation of rhesus macaque-reactive antibody reagents. Representative chromogenic immunohistochemistry images for all antibody targets in spleen, lymph node, lung, or gastrointestinal (GI) tissue.

**Figure S3:**
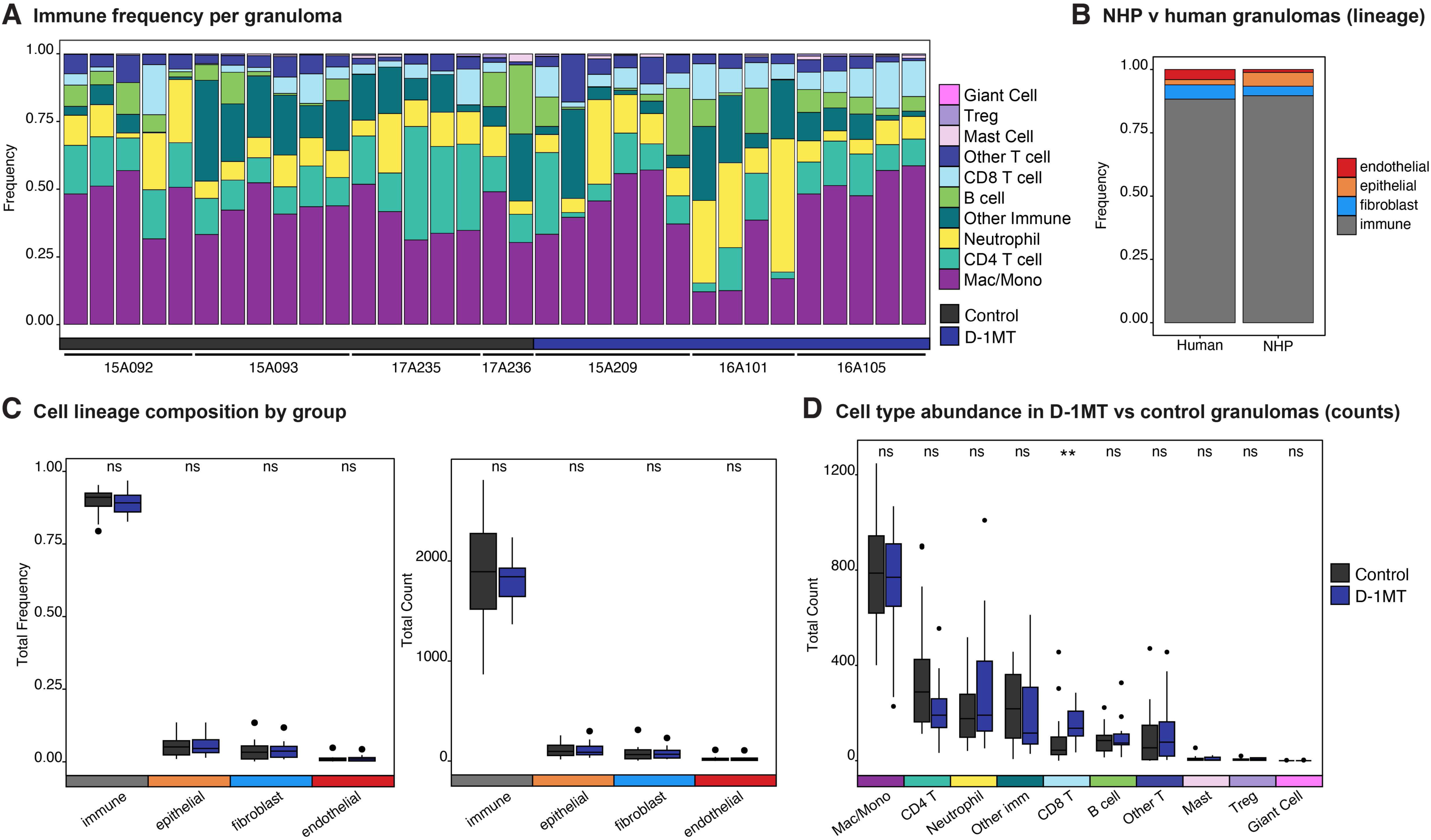
Cellular composition of study samples. **(A)** Proportion of each immune cell subset (of all immune cells) per acquired FOV and categorized by RM and treatment group. Cell subsets are ordered by median frequency. **(B)** Proportion of major lineages (endothelial, epithelial, fibroblast, and immune) across of RM FOVs in comparison to those from human samples (McCaffrey et. al., 2022). Major lineages are order alphabetically, in descending order. **(C)** Frequency (left) and total counts (right) distribution comparison of each major lineage between granulomas between control and D-1MT-treated RMs. **(D)** Total counts distribution comparison of each immune cell subset (of all immune cells) between granulomas from D-1MT-treated RMs and control RMs. All p-values were calculated with a Wilcoxon Rank Sum test (ns p > 0.05, **p < 0.01).

**Figure S4:**
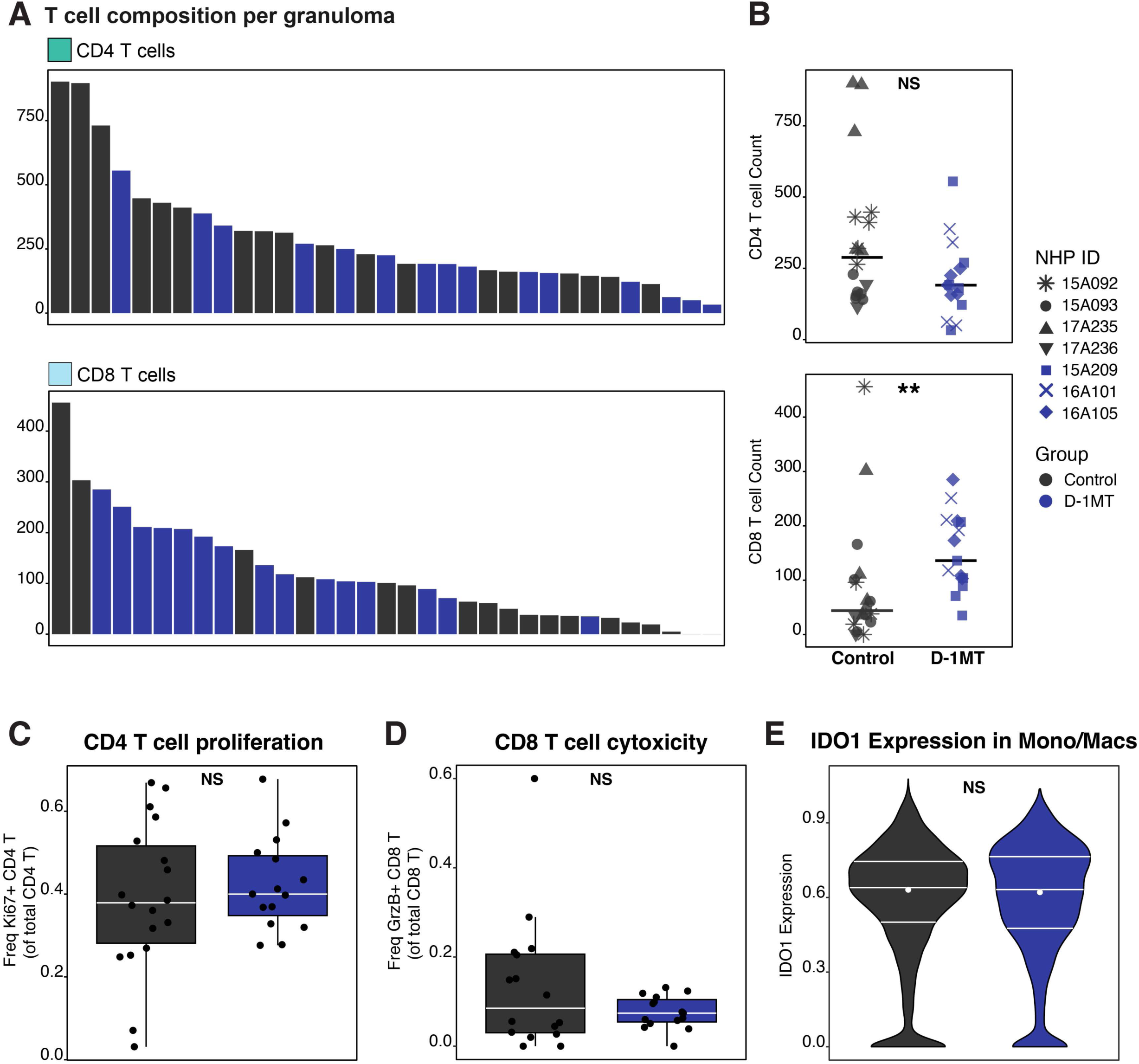
T cell prevalence and immune cell functional status. **(A)** Total counts of CD4+ T cells (top) and CD8+ T cells (bottom) for all FOVs, colored by treatment group and ordered in descending order. **(B)** Quantification of CD4+ T cell (top) and CD8+ T cell (bottom) total counts, stratified by FOVs of granulomas from control RMs and D-1MT-treated RMs. **(C)** Frequency of Ki67+ CD4+ T cells among control and D1-MT-treated RMs. **(D)** Frequency of GranzymeB (GrzB)+ CD8+ T cells among control and D1-MT-treated RMs. **(E)** Violin plots denoting IDO1 expression among macrophage/monocytes, stratified by treatment group. All p-values were calculated with a Wilcoxon Rank Sum test (ns p > 0.05, **p < 0.01).

**Figure S5:**
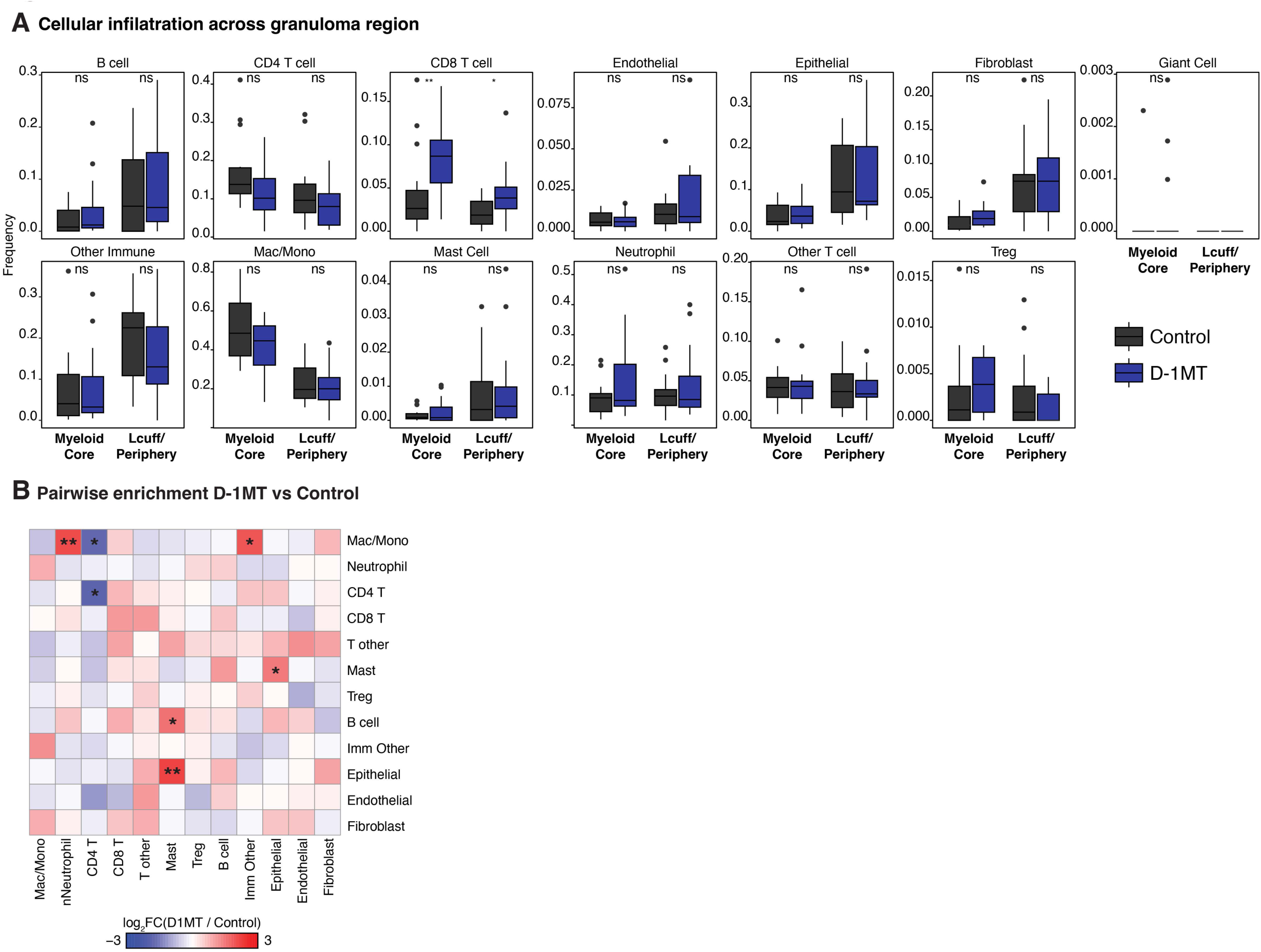
Spatial enrichment of immune cell subsets within RM granulomas. **(A)** Quantification of the local frequency of each immune cell subset (of total cells) in the myeloid core or lymphocytic cuff (Lcuff)/periphery, also stratified by treatment group. P-values were calculated with a Wilcoxon Rank Sum test (ns p > 0.05, **p < 0.01). **(B)** Pairwise cell subtype enrichment (or depletion), shown as a heatmap, considering the effect of D-1MT treatment by taking the log2 fold change of D-1MT and control FOVs. P-values were calculated with a Wilcoxon Rank Sum test (*p < 0.05, **p < 0.01).

