## Supplementary material for "Therapeutic remodeling of the tuberculosis granuloma with 1-methyl-D-tryptophan enhances CD8^+^ T cell-macrophage interactions": All supplemental tables

| Day 1 (overnight stain) | | | | | | | | Parameters for Analysis | |
|---|---|---|---|---|---|---|---|---|---|
| Antibody Target | Biological Relevance | Provider | Catalog Number | Lot | Clone | Mass Channel | Titer (μg/mL) | Start | Stop |
| Chymase | Mast cells | Abcam | ab233729 | GR32553931 | EPR13136 | 71Ga | 0.25 | 70.7 | 71 |
| Tryptase | Mast cells | Abcam | ab181724 | GR32471283 | EPR9522 | 71Ga | 0.25 | 70.7 | 71 |
| CD40 | Costimulation | Abcam | ab13545 | GR322200511 | Polyclonal | 142Nd | 1.00 | 141.7 | 142 |
| CD4 | T cells | IonPath | 714301-100 | 19147-03 | EPR6855 | 143Nd | 1.00 | 142.7 | 143 |
| CD14 | Monocytes/Macrophages | Cell Signaling Technology | 56082BF | 2 | D7A2T | 144Nd | 0.50 | 143.7 | 144 |
| Foxp3 | Regulatory T cells (Tregs) | BD Biosciences | 624084 | 8353822 | 236A/E7 | 146Nd | 1.00 | 145.7 | 146 |
| CD31 | Endothelium | Abcam | ab216459 | GR32965961 | JC/70A | 148Nd | 0.50 | 147.7 | 148 |
| GrzB | Cytotoxic cells | IonPath | 715002-100 | 19235-13 | D6E9W | 150Nd | 1.00 | 149.7 | 150 |
| CD209 | Myeloid-lineage cells | BD Biosciences | 624084 | 9178683 | DCN46 | 152Sm | 0.13 | 151.7 | 152 |
| CD206 | Anti-inflammatory (M2-like) Macrophages | R&D Systems | MAB25341 | CEZR0219091 | 685645 | 153Eu | 0.50 | 152.7 | 153 |
| pS6 | Activation | Cell Signaling Technology | 4858BF | 20 | D57.2.2E | 154Sm | 0.25 | 153.7 | 154 |
| CD68 | Macrophages | Cell Signaling Technology | 76437BF | 2 | D4B9C | 156Gd | 0.13 | 155.7 | 156 |
| CD11c | Myeloid-lineage cells | Abcam | ab212508 | GR32258196 | ITGAX/1242 | 157Gd | 0.25 | 156.7 | 157 |
| CD8 | T cells | Cell Signaling Technology | 85336BF | 4 | D8A8Y | 158Gd | 0.25 | 157.7 | 158 |
| CD3ε | T cells | Cell Signaling Technology | 8506BF | 4 | D7A6E | 159Tb | 0.50 | 158.7 | 159 |
| IDO1 | Immunoregulation | Abcam | ab245737 | GR32593452 | SP260 | 160Gd | 0.50 | 159.7 | 160 |
| CD163 | Anti-inflammatory (M2-like) Macrophages | Cell Signaling Technology | 93498BF | 2 | D6U1J | 163Dy | 2.00 | 162.7 | 163 |
| CD20 | B cells | Cell Marque | 120M-8-OEm | 1908402 | L26 | 164Dy | 0.50 | 163.7 | 164 |
| CD16 | FcγRIII, Myeloid cells and Natural Killer cells | Cell Signaling Technology | 24326BF | 2 | D1N9L | 165Ho | 1.00 | 164.7 | 165 |
| IFNγ | Inflammatory cytokine | Abcam | ab218890 | GR32748436 | IFNG/466 | 166Er | 1.00 | 165.7 | 166 |
| HLA-DR | MHC class II | Abcam | ab7856 | GR32955292 | CDR/43 | 167Er | 0.50 | 166.7 | 167 |
| CD11b | Myeloid-lineage cells | Abcam | ab187537 | GR3245661-3 | EP1345Y | 168Er | 0.25 | 167.7 | 168 |
| CD45 | Immune cells | Cell Signaling Technology | 13917BF | 7 | D9M8I | 169Tm | 0.50 | 168.7 | 169 |
| H3K9Ac | Chromatin accessibility | Cell Signaling Technology | 9649BF | 12 | C5B11 | 170Yb | 0.50 | 169.7 | 170 |
| PanCK | Epithelium | ThermoFisher | MS-343-PABX | 19.2FZ343X1810A | AE1/AE3 | 171Yb | 1.00 | 170.7 | 171 |
| H3K27me3 | Chromatin accessibility | Cell Signaling Technology | 9733BF | 11 | C36B11 | 172Yb | 0.25 | 171.7 | 172 |
| Calprotectin | Neutrophils | ThermoFisher | MA1-81381 | UE2778733 | MAC387 | 174Yb | 0.75 | 173.7 | 174 |
| Ki67 | Proliferation | Cell Signaling Technology | 9449BF | 11 | 8D5 | 175Lu | 0.25 | 174.7 | 175 |

| Day 2 (1 hour) | | | | | | | | Parameters for Analysis | |
|---|---|---|---|---|---|---|---|---|---|
| Antibody Target | | Provider | Catalog Number | Lot | Clone | Mass Channel | Titer (μg/mL) | Start | Stop |
| HH3 | Nuclei | Cell Signaling Technology | 4499BF | 15 | D1H2 | 89Y | 2.00 | 88.7 | 89 |
| Vimentin | Cytoplasm | Cell Signaling Technology | 5741BF | 7 | D21H3 | 113In | 2.00 | 112.7 | 113 |
| SMA | Myofibroblasts | Cell Signaling Technology | 19245BF | 4 | D4K9N | 115In | 2.00 | 114.7 | 115 |
